# Critical Role of Insertion Preference for Invasion Trajectory of Transposons

**DOI:** 10.1101/2022.04.11.487916

**Authors:** Manisha Munasinghe, Nathan M Springer, Yaniv J Brandvain

## Abstract

Transposable elements (TEs) are mobile DNA sequences that have been highly successful at invading eukaryotic genomes. It is unclear how TE families reach high copy number given the expectation that some novel insertions will be deleterious. It has been hypothesized that TE families may evolve to target and insert into specific DNA sequences to adjust the underlying distribution of fitness effects for new insertions. Preferentially inserting into neutral sites could minimize the cumulative deleterious load of a TE family, allowing the mean TE copy number to increase with less risk for host population extinction. To test this hypothesis, we constructed simulations to explore how the transposition probability and insertion preference of a TE family influence the evolution of mean TE copy number and host population size, allowing for extinction. We find that extinction is most common in our simulations under high transposition probabilities, but, as we reduce transposition rates, the risk of extinction persists while the preference for neutral insertion sites is high. In the absence of mechanisms that regulate TE transposition, a preference for neutral insertion sites is not protective and, in fact, actively accelerates both an increase in TE copy number and the time to population extinction.

## Introduction

Transposable elements (TEs) are mobile repetitive DNA sequences that actively increase their copy number and propagate themselves within genomes. A preeminent example of selfish DNA, TEs have been highly successful at invading eukaryotic genomes (Wicker et al. 2007). TEs likely invade naïve populations via horizontal transfer, where the TE moves into the germline of the recipient population and then spreads throughout the genome as well as the population via vertical transmission (Le Rouzic and Capy 2005). TEs employ either a copy-and-paste or cut-and-paste mechanism to insert into novel positions and increase their mean copy number within a population. Class I elements, or retrotransposons, use an RNA intermediate that is reverse-transcribed and integrated into a new position in the genome, while Class II elements, or DNA transposons, move via a DNA intermediate (Feschotte and Pritham 2007; Craig et al. 2015). Insertions that can transpose on their own, as they encode the proteins necessary for transposition, are considered autonomous elements, while non-autonomous elements lack these sequences and consequently rely on autonomous TEs of the same type in order to transpose (Feschotte et al. 2002; Wessler 2006). Classes consist of both autonomous and non-autonomous elements and can be further divided into subclasses, superfamilies, and families depending on their ancestral origins, sequence similarity, and insertional preferences highlighting the genetic and mechanistic diversity of TEs (Kapitonov and Jurka 2008; Seberg and Petersen 2009; Arkhipova 2017).

TE abundance varies greatly between species and is correlated with genome size (Kidwell 2002; Wells and Feschotte 2020). The proportion of the genome occupied by TEs ranges from around 10% in *Arabidopsis thaliana* (The Arabidopsis Genome Initiative 2000), 20% in *Drosophila melanogaster* (Quesneville et al. 2005), and 85% in *Zea mays* ssp. *mays* (Schnable et al. 2009). It is unclear what factors determine not only how much of the genome is occupied by TEs but also how that proportion is distributed between the distinct TE families. Some TE families contribute relatively little to this overall proportion, with only a handful to tens of copies present in a genome, while others contain tens of thousands of copies (Sutton et al. 1984; Baucom et al. 2009; Diez et al. 2014; Stitzer et al. 2021). High TE abundance is surprising given the expectation that novel TE insertions are likely to be deleterious, as they may insert into functional genes, alter heterochromatin formation and gene expression patterns, and induce large structural changes via ectopic recombination (Hedges and Deininger 2007; Hollister and Gaut 2009; Lee and Langley 2012; Adrion et al. 2017).

In the 1980s, Charlesworth and colleagues developed a series of theoretical population genetic models to explore how the mean copy number per individual, commonly denoted by 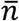, for a given TE family could change over time given their potentially deleterious effects. These studies demonstrated that the mean copy number could increase and eventually stabilize at an equilibrium point depending on the extent of transposition and excision, the effective population size of the host population, and the strength of selection against high TE copy numbers in individuals (Charlesworth and Charlesworth 1983; Charlesworth and Langley 1986; see Charlesworth et al. 1994 for a summary of the effects of these forces). An equilibrium is reached when either all occupiable sites are filled with TEs or the transposition rate is scaled down such that the number of new TEs each generation matches the number lost to drift, selection, and excision. Many of these models make several notable assumptions including: an inverse relationship between transposition and copy number, which limits the number of new insertions as the copy number grows as a built in mechanism of TE regulation, a fixed or increasing fitness effect for new insertions, and a fixed or infinite population size. While these assumptions are quite standard, this means that we have not fully explored the consequences of uncontrolled TE proliferation and how key characteristics of TE families influence their copy number over time.

One aspect of TE biology that has largely been overlooked by population genetic models is the tendency of TE families to preferentially insert into specific DNA sequences or features. Insertion preference is, however, an increasingly important aspect of TE biology as TEs exhibit specific insertion preferences in not only a family manner but also a host dependent manner. The P element in *Drosophila melanogaster* shows strong insertional preference for GC-rich regions near gene promoters, while retrotransposons in several species target sequences upstream or downstream of tRNA genes (Liao et al. 2000; Blanc and Adams 2004; Spradling et al. 2011; Asif-Laidin et al. 2020 p.). The Ty5 retrotransposon in *S. cerevisiae* preferentially inserts into heterochromatin found at the telomeres, which would silence further transposition of the novel insertion but also avoids any deleterious effect associated with inserting into a gene (Boeke and Devine 1998; Gao et al. 2008; Novikova 2009). Consequently, insertion preference may not only be the result of structural differences between TEs but also an evolved trait that impacts the expected selective effect of new insertions. The genetic load of TE families that preferentially insert into heterochromatic or intronic regions is expected to be less deleterious than those that insert into functional or genic regions. Insertion preference therefore represents not just the nucleotide sequence or feature a TE family inserts into but also dictates an underlying distribution of fitness effects for each new insertion. To allow for such preference, Charlesworth (1991) developed a deterministic model consisting of two classes of TE insertions, selected against or neutral, and found it difficult to obtain combinations of parameters for transposition, excision, and selection against insertions that matched TE copy numbers observed in *Drosophila* (Charlesworth 1991), suggesting that an insertion preference cannot stabilize TE proliferation. However, this work did not consider the stochastic nature of TE replication nor did this work include the possibility that the genetic load imposed by TEs could decrease population growth rates, ultimately leading to population extinction.

It is obvious that unconstrained TE transposition would cannibalize the host genome and ultimately drive the host population extinct, but we know comparatively little about how variation in TE biology impacts this process. In the absence of mechanisms that manage or eliminate transposition, such as auto-regulation or host repression, a clear expectation is that highly replicative TE families should most rapidly expand in copy number and drive populations extinct. Conversely, TE families that evolve a preference for neutral insertion sites should allow the copy number to increase without growing the deleterious genetic load, potentially allowing the host population to survive for longer. However, since nearly all population genetic models assume either a fixed or infinite host population size, we do not have actual confirmation of these hypotheses or an understanding of how these facets of TE biology influence mean copy number and population size over time.

Here, we use a non-Wright-Fisher framework in SLiM 3 to explore how transposition probability and insertion preference influence the evolution of mean TE copy number and host population size (Haller and Messer 2019). We consider a naïve diploid population that gains a single copy of a TE in the genome of a single individual (analogous to horizontal transfer). This TE belongs to a unique family with an assigned transposition probability and range of fitness effects for novel insertions that represent insertion preference. TEs transpose and increase their mean copy number in the population over time, and we manage transposition such that the probability of any given element copying and inserting itself elsewhere in the genome is held constant to mimic an unregulated proliferation. If the TE family is not initially lost due to either genetic drift or selection, we can track its spread through the population by measuring the mean copy number and population frequency of the TE family. We allow population size to fluctuate depending on the fitness of individuals, such that populations can go extinct. Consequently, we can relate changes in the mean copy number to the mean fitness of the population to explore how key aspects of TE biology influence the invasion trajectories of TE families and under what conditions populations survive the invasion.

## Methods

### Model Setup

We consider a diploid, hermaphroditic population that reproduces sexually in SLiM 3 (v3.6). At the beginning of a simulation, we introduce a single TE in the genome of a single individual. This TE belongs to a unique family with a specified transposition probability (*teJumpP*) and preference for either neutral or deleterious insertions (*neutP*). We can track the mean copy number of the TE family in an individual over time along with changes to the mean fitness and size of the population. Consequently, we can assess under what conditions the copy number increases and whether that increase could drive the population extinct. Our model design is based on recipe 14.12 (Modeling transposable elements) in the SLiM manual (Haller and Messer 2016) with a few extensions. Notably, we extend this model to employ a non-Wright-Fisher (nonWF) framework, which allows the population size to fluctuate depending on the fitness of individuals, and incorporate insertion preference, which allows TE insertions to have variable fitness effects. We comment on key aspects of our model below, and full details of our model can be found in the supplement (ESM Appendix 1).

### Modeling Genome Architecture

Previous population genetic theory has demonstrated the influence of recombination rate on TE accumulation patterns as a result of Hill-Robertson effects and Muller’s ratchet (Muller 1964; Hill and Robertson 1966; Langley et al. 1988; Dolgin and Charlesworth 2008). We consequently consider one genomic architecture with low recombination and one with higher recombination. We first consider a genome consisting of a single chromosome (*L* = 1 x 10^5^ occupiable sites) with a uniform recombination rate of *r* = 1x 10^-8^. We then considered an architecture modeling 5 distinct chromosomes (*L* = 5 x 10^5^ occupiable sites divided equally) with a uniform recombination rate of *r* = 1 x 10^-5^. It is worth clarifying that we do not model the actual nucleotide sequence of TEs or the host genome. Each position in the genome represents an occupiable site with an associated fitness effect if occupied by a TE. New insertions do not change the length of the genome, and we do not rack the connections between each new insertion (i.e., which TE a novel TE derived from).

### Modeling TE Insertion Preference

Insertion preference in our model is not a specific sequence or feature that a TE inserts into but instead a representation of an underlying distribution of fitness effects for novel insertions. It effectively adjusts the likelihood that a novel insertion will be neutral versus deleterious. For simplicity, we pre-assign each site in the genome to one of four selective classes that represent the fitness effect to an individual if a TE inserts into that site: neutral (*s* = 0.0), mildly deleterious (*s* = −0.005), modestly deleterious (*s* = −0.05), and massively deleterious (*s* = −0.5). The proportion of each class in the genome is determined by the parameter *neutP* which represents the probability that a site in the genome is neutral (i.e., the number of neutral sites in the genome for a given parameter set is equal to *L* * *neutP* where *L* represents the number of sites in the genome) with the remaining sites equally divided between the 3 different deleterious classes. Each position assumes a dominance coefficient of *h* = 0.5 such that the full fitness effect is only realized in individuals homozygous for the TE insertion at that site. Individual final fitness is then calculated multiplicatively across all loci. The positions of the sites for each class are randomly drawn so that there is no artificial clustering of fitness effects and are kept constant across replicates. Consequently, low *neutP* values represent TE families where new insertions are more likely to be deleterious, as the genome consists of more sites that will result in deleterious effects if a TE inserts into them.

### Modeling TE Transposition

Transposition is modeled as a Poisson process with the parameter *teJumpP* representing the probability that a single TE will copy and insert itself elsewhere in the genome. The number of novel insertions for a single individual is then drawn from a Poisson distribution dependent on both *teJumpP* and the number of autonomous elements present in the genome. The site for each new insertion is randomly chosen, and the fitness effect of the novel insertion is then determined based on the assigned fitness consequence if a TE inserts into that site (as detailed in the previous section). *teJumpP* is fixed for the entirety of a simulation run meaning we do not rescale or limit transposition. High *teJumpP* values therefore represent rapidly proliferating TE families.

### Modeling TE Biology

Outside of transposition and insertion preference (detailed above), we consider two notable features of TE biology. As mentioned above, TEs are considered autonomous if they encode the necessary elements required for their transposition; however, it is possible that they may lose features during transposition resulting in non-autonomous elements that can no longer independently transpose. These non-autonomous elements contribute to total copy number and affect the fitness of the host individual, but they rely on present autonomous elements in order to generate any additional insertions. We do extend our models to include non-autonomous elements by assuming that 50% of all novel insertions result in a non-autonomous element. These elements contribute to copy number and fitness, but they are not counted when determining the number of novel insertions that occur. DNA transposons which employ a cut-and-paste mechanism are first excised from one location and then reintegrated elsewhere in the genome. Reintegration can, however, fail resulting in a decrease to copy number as the element is lost (Xu et al. 2004). This random excision not only reduces TE copy number but also increases the chance that the TE family could be lost from the population entirely. We extend our models to include random excision at 1/10^th^ the chance of transposition and randomly determine which TEs are lost from the population.

### Life Cycle, Population Growth, and Selection

We initialize the population with *K* = 1000 individuals, with *K* acting as the hard carrying capacity for the population. The generational life cycle in non-Wright-Fisher models in SLiM starts with the creation of offspring. Each generation, we generate a population that is twice the size of the previous generation (N_t+1_ = 2N_t_) selecting all parents independently and at random. We then employ viability selection such that an offspring’s survival probability is simply its expected fitness – assuming additivity within and multiplicative fitness across loci (see above). We enforce discrete generations in the nonWF SLiM framework by setting the fitness of all parents to 0 before viability selection occurs. Finally, if the number of surviving offspring exceed the population’s carrying capacity, K, we randomly cull excess offspring to generate a population of size no greater than K. Transposition then occurs in the remaining individuals who survived viability selection. Since fitness is evaluated only during viability selection, new or lost elements do not influence the probability of an individual being chosen as a parent in the next generation. They will, however, be inherited by any potential offspring, and, consequently, influence their offspring’s ability to survive. We then loop back to the start of the generational cycle with the surviving offspring forming the new parental pool.

### Model Outcomes

For each distinct parameter combination, we track the mean fitness and size of the population as well as the mean copy number of the TE family and mean frequency of a TE in the family stratified by their fitness effects over time. We consider 3 endpoints for a replicate simulation run of a given parameter set.

#### Outcome 1, TE Loss

Loss of the TE family from the host population occurs if the mean copy number of autonomous TEs is zero. No further transposition can occur, and we simply track which generation the TE family was lost in.

#### Outcome 2, Population Extinction

If the population size hits zero, we record that replicate as resulting in a population extinction event and output the relevant trajectories (population fitness, size, mean TE copy number, and mean TE frequency over time). population extinction occurs when no individuals in the population survive after viability selection.

#### *Outcome 3*, Dual Survival

If neither of these options occur, then both the TE family and the host population have survived to the final generation (capped at 50,000). We output the relevant trajectories (same as though outputted in population extinction) for that simulation run.

### Models, Parameters, and Replicates

We consider 5 distinct models that varied either genome architecture or specific aspects of TE biology detailed above (Table 1). When describing our results, we use the term TE family to refer to a specific parameter combination of transposition probability (*teJumpP*) and insertion preference (*neutP*). We explore the following sets of values for our parameters: *teJumpP* = [1 x 10^-4^, 2.5 x 10^-4^, 5 x 10^-4^, 7.5 x 10^-4^, 1 x 10^-3^, 2.5 x 10^-3^, 5 x 10^-3^, 7.5 x 10^-3^, 1 x 10^-2^, 2.5 x 10^-2^, 5 x 10^-2^, 7.5 x 10^-2^, 1 x 10^-1^] (13 values) and *neutP* = [0.010, 0.025, 0.050, 0.075, 0.10, 0.25, 0.50, 0.75, 0.90, 0.925, 0.950, 0.975, 0.99] (13 values) for each model, resulting in a total of 169 distinct parameter combinations per model type. Each distinct parameter combination is used for a specific simulation run of a model. We initialize the host genome, TE family, and population and consider 3 endpoints (TE loss, population extinction, or dual survival) for a replicate of a given parameter set. When one of these outcomes occur, we record the relevant trajectories and end state before looping back to the initialized state. The simulation run is finally complete (i.e., we no longer loop back to the beginning point) if we record either a combined total of 100 population extinction and dual survival events or 1 x 10^6^ TE loss events.

**Table 1.**
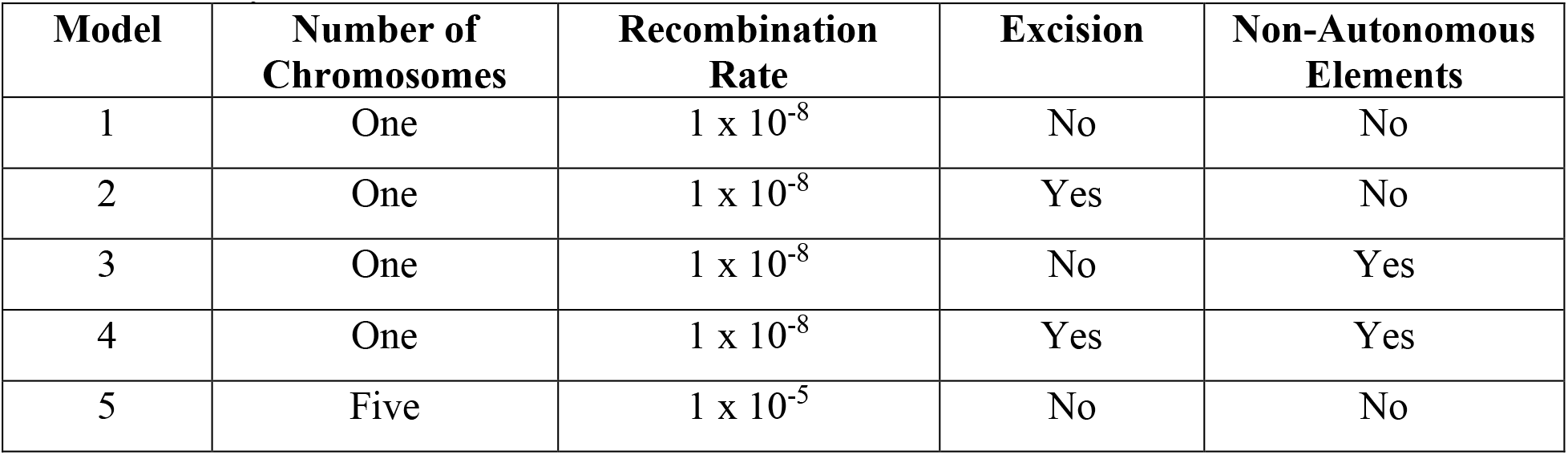
Summary of Models Constructed.

Certain parameter combinations resulted in dramatically longer run times, due to SLiM executing operations over many TEs for many generations. For these combinations, we scaled down the outcome by an order of magnitude (i.e., 10 population extinction and dual survival outcomes to 1 x 10^5^ TE Loss outcomes) and parallelized runs if necessary. In all figures presented, we mark these combinations to highlight this change. Scripts for all models can be found on GitHub – see Data Accessibility statement.

## Results

A set of models were developed to assess the outcome of introducing a TE, which can replicate freely, into a naïve population (example showing our five chromosome model in Figure 1). There are three classes of outcomes for this TE introduction: TE loss, population extinction, and dual survival. TE loss occurs whenever there are no autonomous copies of the TE family remaining in the population, which can occur via drift or selection. Population extinction results when mean fitness is reduced such that no host individuals remain after viability selection. Dual survival refers to instances in which the population survives and retains even a single copy of the TE family for 50,000 generations.

**Figure 1.**
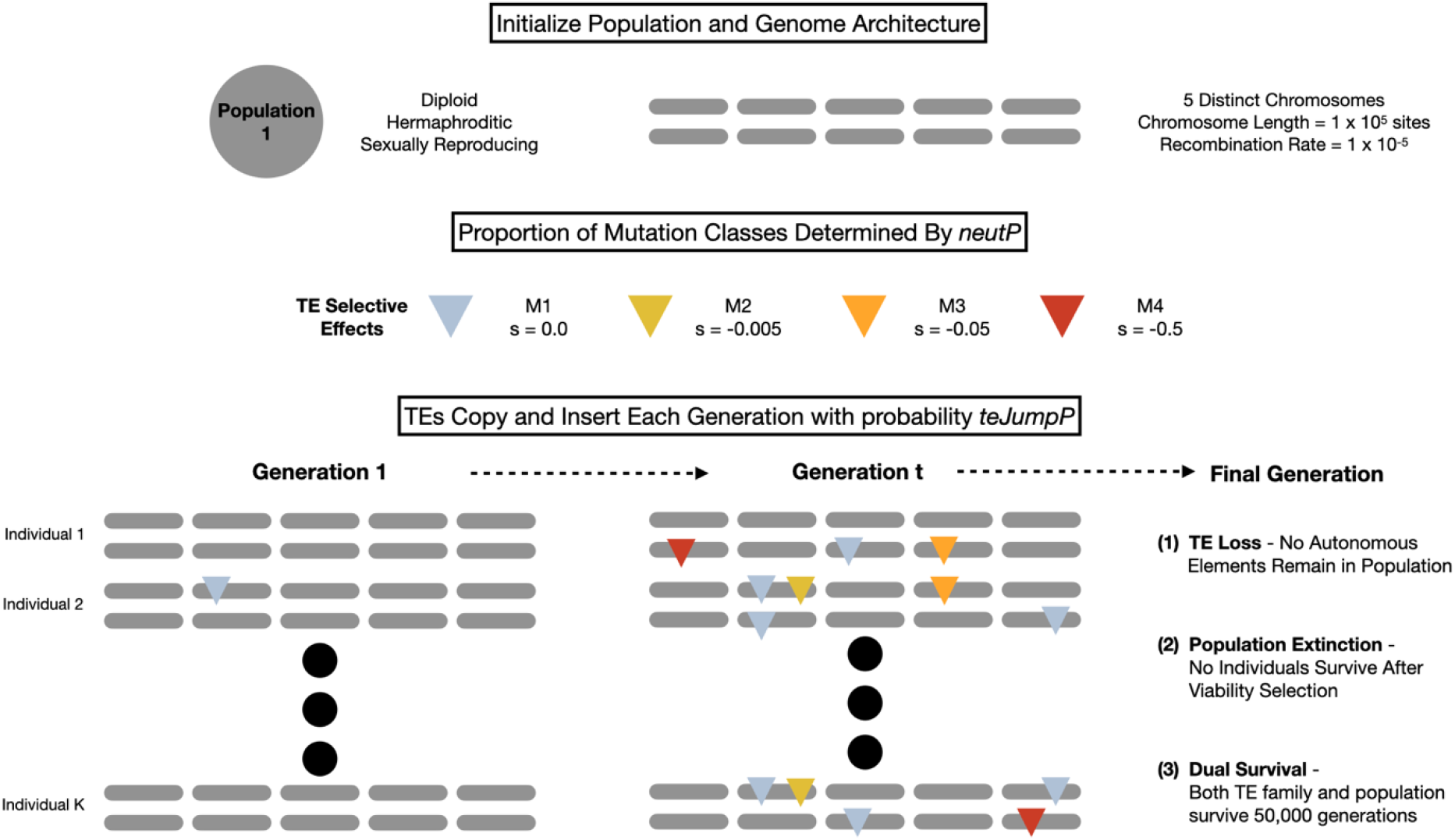
Visual Summary of Five Chromosome Model. A visual representation of the five chromosome model design detailed that highlights the population initialization and the role of the two key parameters, *teJumpP* and *neutP*.

Initially, we explored the proportion of these three classes of outcomes across the parameter space for transposition probabilities and insertion preferences for all five models. Neither the inclusion of excision or non-autonomous elements substantially alters the general trends we observe across our parameter space, and we, therefore, relegate most of the discussion and visualization of those models (Model 2,3, and 4) to the supplement (Figure S1,2). We focus here on the differences between our first model, which contained only one chromosome with relatively low recombination, to that of our last model, which contained five distinct chromosomes with higher recombination.

Before proceeding, we invite the reader to predict which outcomes may prevail across the parameter space. We had several initial expectations about the likely outcomes. One expectation is that a preference for neutral insertion sites will protect the host population by minimizing the cumulative deleterious load of the TE family, allowing dual survival instead of population extinction. Another is that the highest observed TE copy numbers will occur under the highest transposition probabilities. Combined, we predicted that a TE family with a high transposition probability and strong preference for neutral insertion sites may be the most effective at reaching high copy numbers without rendering the host population extinct.

### TE Loss is the Predominant Outcome

By far, the predominant outcome across all parameters is TE loss, which is not surprising given the very low initial allele frequency. Within all replicates for a given parameter combination across all models, at least 84% of the simulations result in TE loss (Table S2). Approximately, 95% of all TE loss outcomes occur within 50 generations, which we expect is primarily due to genetic drift as the initial TE is neutral and begins with frequency 1/2000 (0.0005). In our single chromosome model, TE loss is the sole outcome when the transposition probability is moderate to high and the neutral insertion preference is low (Figure 2A).

**Figure 2.**
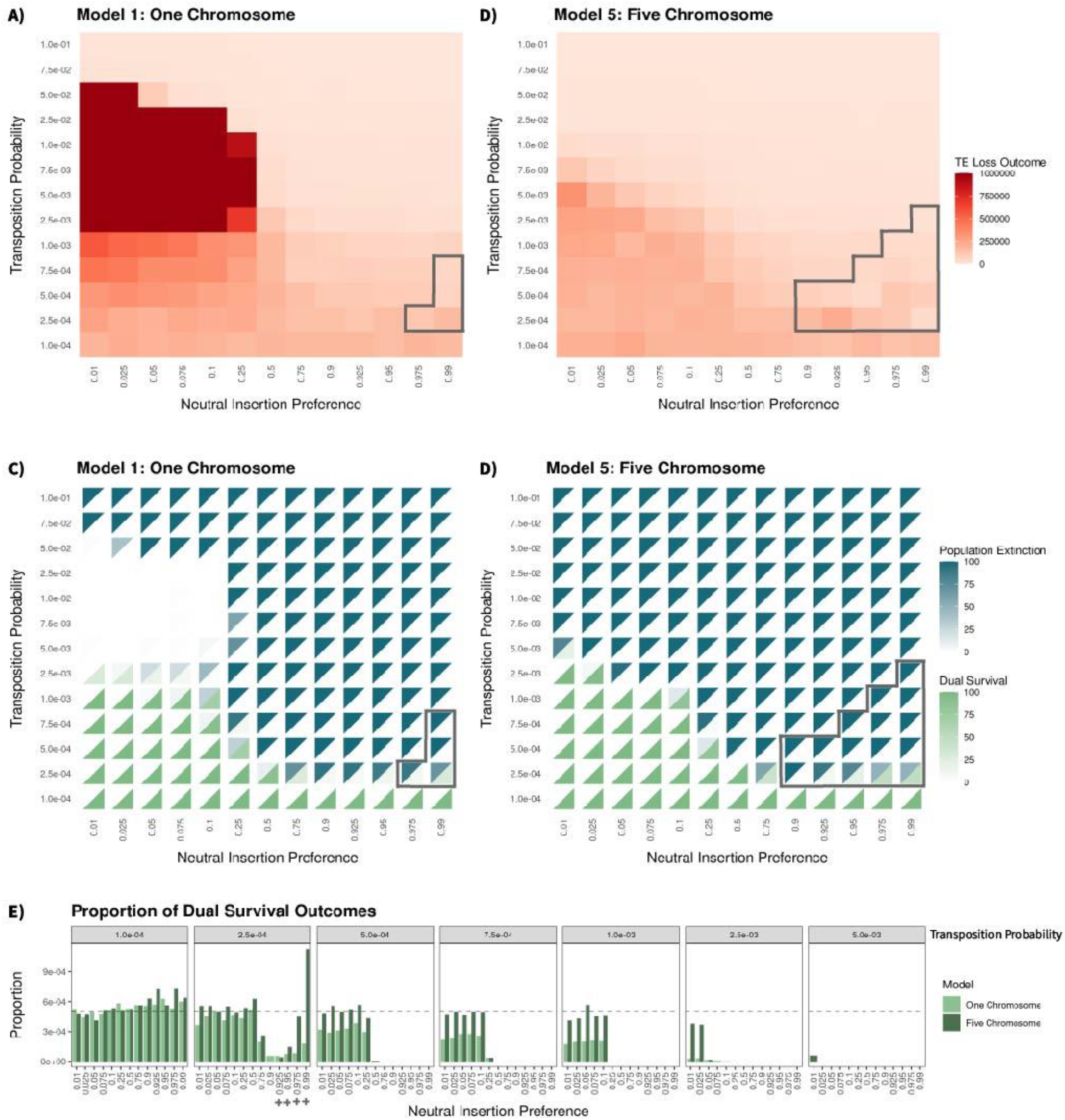
Outcome Across Our Parameter Space. The left column visualizes outcomes for our single chromosome model with *r* = 1 x 10-8, and the right column shows outcomes for our five chromosome model with *r* = 1 x 10-5. The top row (A and B) shows heatmaps colored with the number of TE introductions that resulted in loss of the TE family. The bottom row (C and D) shows heatmaps colored with both the number of TE introductions that resulted in population extinction (upper triangle in blue) or dual survival (lower triangle in green). For each heatmap, rows represent the transposition probability with increasing probabilities as you move upwards, and columns represent the neutral insertion preference with increasing preferences as you move to the right. The final row (E) highlights the proportion of dual survival outcomes compared between models. Cells indicated with grey markers represent parameter combinations that, due to computational constraints, had reduced limits detailed in the methods.

For these combinations, 1 million TE introductions all result in TE loss without a single observed population extinction or dual survival event. This is likely due to background selection against deleterious TEs that, due to the low recombination rate, are linked to neutral TEs which are inadvertently lost. Both excision and the inclusion of non-autonomous elements increase the proportion of TE loss outcomes and expand this region of exclusive TE loss (Figure S1A-B). Excision widens this region to include both higher and lower transposition probabilities, while non-autonomous elements shift the region upwards to include higher transposition probabilities. TE loss is likely the result of both genetic drift and selection. This is further exacerbated under low recombination rates where neutral TEs are strongly linked to deleterious TEs and consequently experience negative selection, purging all TEs from the population.

Our single chromosome model with low recombination demonstrates a higher proportion, sometimes even exclusively, of TE loss (Figure 2A) in comparison with our five chromosome model with higher recombination (Figure 2B). Higher recombination allows neutral TEs to recombine away from deleterious TEs and persist in the population while selection acts on deleterious copies.

The probability of fixation for any neutral mutation is equivalent to its allele frequency. We find that, for the most part, the proportion of TE losses aligns with drift expectations (Fig 2E). The most notable deviations from this neutral expectation occur when we increase the transposition probability, due to an increase in the proportion of population extinction events. If we fix the transposition probability, we find additional increases in population extinction outcomes as we increase the preference for neutral insertion sites.

### TE Family Dynamics Influence Trajectories of Extinction and Survival

The relatively rare TE introductions that do not result in TE loss can be divided into population extinction or dual survival outcomes. The proportion of these outcomes were visualized across the full parameter space of TE transposition probability and insertion preference (Figure 2C,D). Transposition probability is the primary factor in determining whether population extinction or dual survival outcomes occur. Population extinction occurs predominantly when the transposition probability is high, while dual survival occurs when this probability is low. The magnitude of these two outcomes is quite distinct with moderate levels of population extinction occurring under some parameters (up to ~15% of all outcomes) but only very low levels of dual survival (up to ~0.11%). If we fix the transposition probability, we find that the proportion of population extinction outcomes increases as we increase the neutral insertion preference (i.e., more extinction occurs under higher neutral insertion preferences).

We remind the reader that the proportion of outcomes that are observed in our models depends on the maximum number of generations simulated (50,000). The vast majority of TE loss events occur after very few generations. In contrast, extinction events occur over a much broader time gradient with many happening after 10,000) generations. Fitness trajectories for dual survival outcomes indicate that some would have gone extinct if we had extended the simulations for more generations (Figure S3). This highlights the dependency of the proportion of extinction and dual survival events on the generational time limit. We hypothesize that, in the absence of a generational time limit, many of our dual survival outcomes would transition to population extinction events. Extinction, in our model, depends on the deleterious load incurred by the TE family. If TE families are given additional time to proliferate freely, the deleterious load is similarly allowed to grow, ultimately increasing the chance of extinction.

### High Neutral Insertion Preference Accelerates Population Extinction

To examine the factors that are associated with population extinction, we examined the timing of extinction events and the copy number of deleterious and neutral TEs in the generation preceding extinction in our five chromosome model (Figure 3). Populations with higher transposition probabilities go extinct more rapidly across all insertion preferences (Figure 3A). For a fixed transposition probability, lower neutral insertion preferences delayed the time to host extinction when compared to higher neutral insertion preferences. However, the time to extinction eventually stabilizes across the higher range of neutral insertion preferences (Figure 3A).

**Figure 3.**
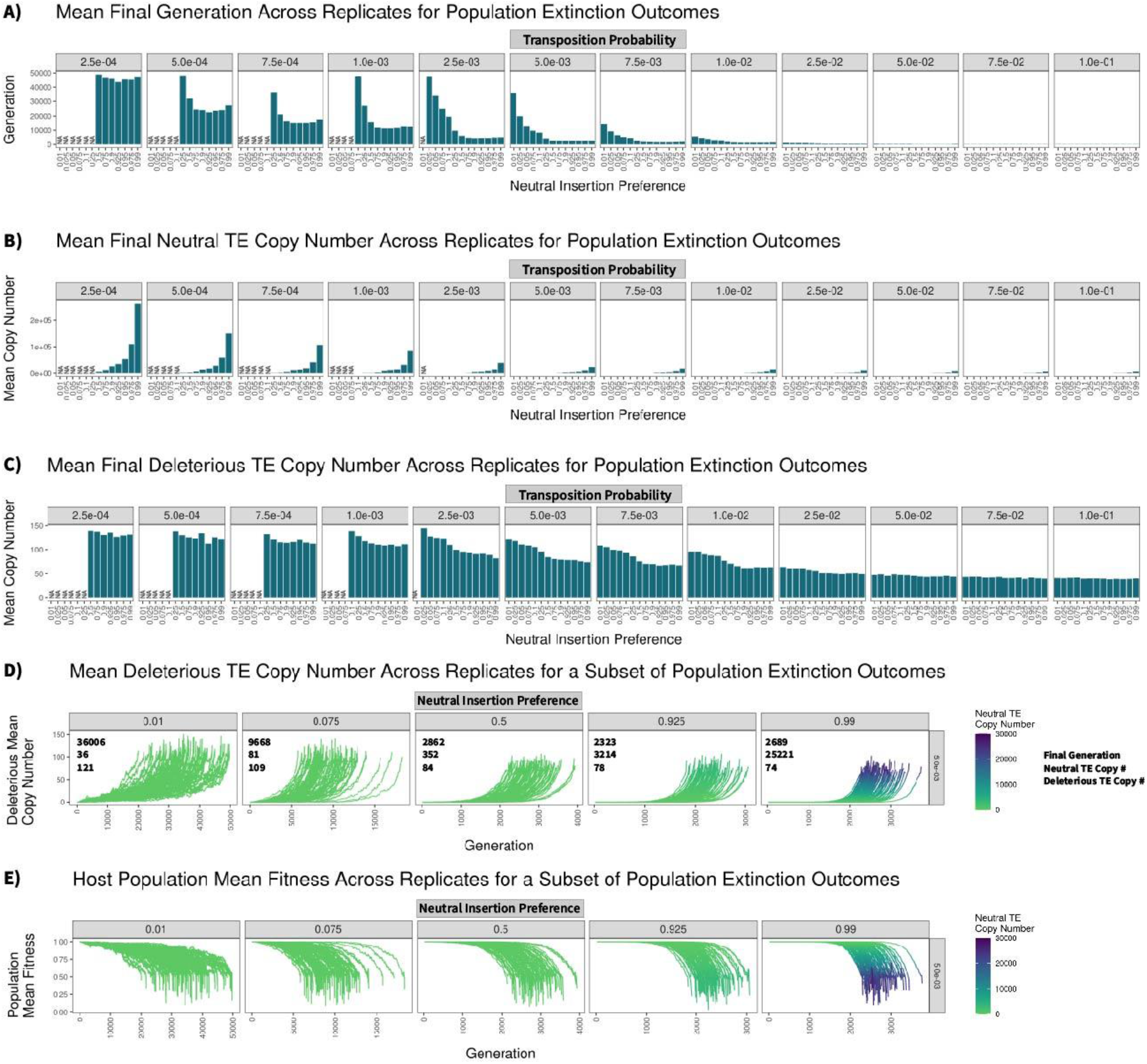
Key Dynamics of Population Extinction Outcomes in the Five Chromosome Model. We compared population extinction outcomes across our parameter space by looking at (A) the mean generation with which the population went extinct, (B) the mean neutral TE copy number in this final generation, and (C) the mean deleterious TE copy number in this final generation across replicates for parameter combinations with observed population extinction outcomes. NAs indicate combinations where no population extinction outcomes were observed. We facet these visualizations by the transposition probability, with greater probabilities on the left, with the x-axis showing increasing neutral insertion preferences. We then fix the transposition probability at *teJumpP* = 5.0 x 10-3 and visualize the change in (D) the deleterious TE copy number and (E) population mean fitness over time for a subset of neutral probabilities for observed extinction outcomes. We facet these graphs by insertion preference and allow the x-axis limits to vary to better visualize not only the trajectories but also how extinction occurs more rapidly under greater neutral insertion preferences.

Population extinction occurs when the deleterious load of the TE family is sufficiently high such that no individuals survive viability selection. The relationship between the final mean copy number of neutral and deleterious TEs in extinction events was monitored (Figure 3B-C). There is high variability for the copy number of the TE family in the population at the final generation before extinction across our parameters. Because populations face steeper fitness declines when transposition is frequent, the final copy number tends to decrease as we increase the transposition probability and reduce the preference for neutral insertions, particularly for neutral TE copy number (Figure 3B). Consequently, the highest final copy numbers are observed under moderate transposition probabilities with a strong preference for neutral insertions. There is substantial variability in the final neutral copy numbers (as few as 0 and as many as 270,000) but relatively little variation in the copy number of deleterious TEs (Figure 3C). We observe higher deleterious copy number numbers at low transposition probabilities (~125 copies) in comparison to high transposition probabilities (~50 copies). While the neutral insertion preference strongly impacts the time to extinction and the neutral TE copy number, it has a very limited impact upon the final deleterious TE copy number.

We find that the copy number of deleterious TEs stays low until a critical threshold of TEs is reached, at which point the deleterious copy number grows exponentially (Figure 3D). Populations enter an extinction vortex at this threshold, with a rapid decline in subsequent fitness driven by an accumulation of deleterious TE insertions. The TE copy number required to trigger this process depends on both the transposition probability and insertion preference. High transposition probabilities require fewer present TEs to generate high numbers of novel insertions, which explains the lower copy numbers observed in populations immediately before extinction under high transposition (Figure 3B,C).

Higher preferences for neutral insertions require more TE copies since a smaller portion of novel insertions in the next generation are expected to be deleterious. Since selection acts on deleterious TE insertions, a strong preference for neutral TE insertions allows populations to reach this critical threshold more rapidly as neutral TEs accumulate and provide a source for higher numbers of insertions in the next generation. This explains the tendency to observe shorter times to extinction under high neutral insertion preferences as well.

We initially hypothesized that a higher preference for neutral insertion sites would protect the host by reducing the deleterious load incurred by insertions. Instead, our models show that higher preferences for neutral insertions increase the risk of population extinction, eliminating both host and TE. While this result may be counterintuitive at first, it makes sense given the design of our model. We do not rescale the transposition probability given the current copy number. For a fixed transposition probability, the number of expected novel insertions for a given TE copy number is the same across insertion preferences. It is the proportion of neutral to deleterious TE insertions that varies with insertion preferences. Higher neutral insertion preferences means the expected proportion of novel insertions that are expected to be neutral is greater, but it does not fully eliminate the chance of observing deleterious insertions. When the number of novel deleterious insertions each generation is small, selection can purge them from the population; however, if the number of deleterious insertions increases each generation at a rate faster than selection can remove them, the population will enter into an extinction vortex as the deleterious copy number grows exponentially. A preference for neutral insertion sites allows TE families to more rapidly reach high copy numbers guaranteeing an increasing number of deleterious TE insertions in subsequent generations. Thus, while higher neutral insertion preferences allows a TE family to rapidly invade and proliferate, it ultimately leads to extinction in scenarios with uncontrolled proliferation where neutral TEs ultimately churn out more deleterious TEs than the population can withstand.

### TE Survival Can Result in Variable Final TE Copy Number and Allele Frequency

While the range of observed final mean copy numbers is comparable across population extinction and dual survival outcomes, the distribution is not.

We exclusively observe very low copy numbers under dual survival, with just over 50% of all observed dual survival outcomes exhibiting a final copy number of less than 10. The smallest copy number observed under population extinction, for comparison, is 21. Similarly, the median final copy number for dual survival outcomes is ~9 as opposed to ~587 in population extinction, highlighting the elevated range of copy number for population extinction. The largest observed copy number is relatively similar between population extinction (268,699) and dual survival (245,171); however, many fitness trajectories for high copy number (> 1000) dual survival outcomes are similar to population extinction trajectories (Fig S3,4). This indicates that these outcomes would have gone extinct if we had extended the generational time limit, which raises doubt to the stability of high copy number dual survival outcomes (however this slow extinction may allow populations enough time to evolve mechanisms to suppress and remove such TEs). The only consistently stable dual survival trajectories occur under our lowest transposition probability (*teJumpP* = 1.0 x 10^-4^) with a maximum copy number of 1,211 across all measured neutral probabilities.

Looking exclusively at dual survival outcomes, we find that the final copy number does increase as we increase the preference for neutral insertion sites; however, many outcomes still exhibit copy numbers fewer than 100 (Fig 4A). When transposition is low, high copy number (>100) dual survival outcomes occur only under high neutral insertion preferences. While high transposition often results in extinction, we do observe dual survival outcomes as long as the neutral insertion preference is reduced. Here, high copy numbers can be observed under low to moderate neutral insertion preferences, but this space can easily transition to population extinction if either parameter is increased. Once a single TE fixes in the population, dual survival will occur as long as the mean copy number of the TE family stays below the critical threshold. When looking at the final allele frequency of a single TE within a replicate, we observe distinctly bimodal distributions (Fig 4B). There are a substantial number of low frequency polymorphic (< 0.25 frequency) TEs and fixed TEs (albeit reduced in comparison to polymorphic TEs), but we observe very few moderate frequency TEs (between 0.25 and 1.0 frequency).

**Figure 4.**
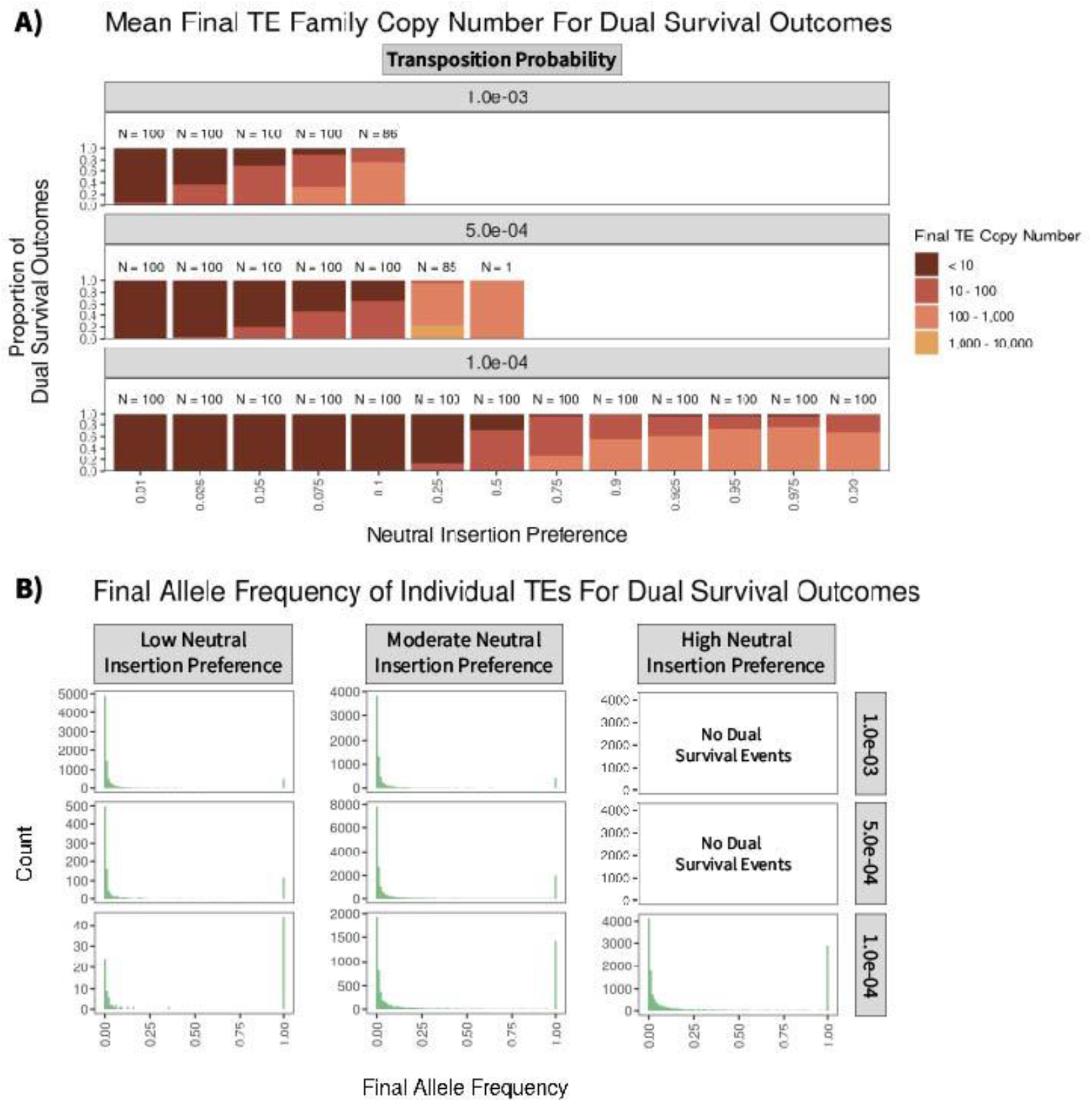
End States Observed for Dual Survival Events in our Five Chromosome Model. (A) Shows the distribution of the mean final TE Copy Number of the host population. We bin the final mean copy number for a given replicate into one of four ranges (color). We show the insertion preference on the x-axis and the number of replicates for each bin on the y-axis for three transposition probabilities that span the range of dual survival outcomes. The number above indicates how many dual survival outcome events were observed. (B) Shows the distribution of final allele frequencies for every specific TE (i.e., for a site in our genome occupied by a TE how many individuals also carry a TE at that site). We bin the neutral insertion preference into either low (0.01 – 0.075), moderate (0.1 – 0.9), or high (0.925 – 0.99) and once again subset three transposition probabilities that span the range of dual survival outcomes. The x-axis of each plot shows the allele frequency, and the y-axis shows how many TEs had that frequency in the final generation.

## Discussion

We construct a series of models to simulate the invasion of a naïve population by a TE family with a unique transposition probability and insertion preference. While loss of the TE family from the population dominates across the entirety of our parameter space, we find that population extinction and dual survival differentially occur in somewhat surprising ways. Strikingly, we find a counterintuitive relationship between insertion preference and population extinction. Population extinction occurs most frequently under high transposition probabilities, but as we reduce transposition probability, we find that population extinction is not only more common when transposition is more biased towards neutral sites but also occurs more rapidly. A preference for neutral insertion sites does result in the highest final TE copy numbers, but these outcomes always result in population extinction. Dual survival occurs only under low transposition and is characterized by notably smaller copy number.

Our results reject the hypothesis that preferentially inserting into neutral sites allows the TE copy number to grow by minimizing the cumulative deleterious effects of insertions, consequently protecting the host population. In the absence of mechanisms that regulate transposition, a preference for neutral insertion sites does allow the mean copy number to grow more rapidly due to the absence of selection on neutral insertions. However, the proportion of expected deleterious insertions remains the same. Once the TE copy number is sufficiently high enough, the raw number of deleterious TE insertions each generation grows faster than selection can remove them. Consequently, the population enters into an extinction vortex driven by an exponential increase in the deleterious TE copy number. Reducing both the transposition probability and preference for neutral insertions results in more dual survival, but this outcome is characterized predominantly by very low mean copy numbers.

Our models consider two distinct host genetic architectures (one chromosome with low recombination or five chromosomes with high recombination) and biological features of TE families (random excision of elements, non-autonomous elements, both, or neither). We find that low recombination results in higher proportions of TE loss, especially when the preference for neutral insertion sites is low. Neutral TEs are tightly linked with deleterious TEs and consequently purged often resulting in loss of the TE family from the population, in a process analogous to Muller’s ratchet (Muller 1964). A higher recombination rate minimized this effect, and we did not observe any regions of exclusive TE loss under this scheme. We find that neither the addition of random excision of TEs nor the inclusion of non-autonomous elements substantially alters our results. These characteristics of TE biology simply influence the amount of TE loss outcomes and shift the region of exclusive TE loss observed. This is expected as both features increase the chance there will be no remaining autonomous TEs in the host population.

Given unregulated transposition, we find that high copy numbers almost always result in population extinction and that even an evolved preference for neutral insertion sites is not enough to avoid this outcome. The assumption that TE families reach high copy number may be linked to their characterization as selfish genetic elements seeking to proliferate regardless of their effects on the host (Dawkins 1976; Orgel and Crick 1980; Werren et al. 1988). However, achieving high copy number at the expense of the host population affects the TE family as well as they are consequently wiped out. Population extinction effectively culls host populations incapable of silencing TE proliferation and the highly-replicative TE lineage. In natural populations, high TE copy number is in fact less common than one may expect. A recent analysis by Stitzer et al. (2021) annotated structurally intact TEs in the maize reference genome to investigate family-level dynamics of TEs in maize. 75% of their observed TE families contain only a single copy, with 95% of all families showing less than 10 copies. Only 1.2% of all families exhibited copy numbers larger than 100, and only four families in total had copy numbers greater than 10,000 (Stitzer et al. 2021). In a similar study in *Arabidopsis*, only four TE families exhibited copy numbers greater than 1,000 (all Helitrons with a max copy number of ~1,500) (Ahmed et al. 2011; Quesneville 2020). This work suggests that many more TE families exist at low copy number than high copy number. Selection obviously contributes to this differential, but, in addition to purging deleterious insertions, there may also be selective pressure for more moderate transposition probabilities as well. This could limit the proliferation of TEs in order to reduce the chance of driving the population extinct. Stritt et al. (2021) recently challenged the idea of TEs as “invasive” genetic elements replicating aggressively in the absence of silencing mechanisms, resulting in “bursts” of TE activity (Stritt et al. 2021). Their analysis of TE copy number in angiosperms also finds that many TE families exist at low copy numbers, and they suggest that TEs may be maintained in evolution not just because of their ability to independently replicate but also because they have evolved strategies to persist at low copy numbers.

Charlesworth (1991) recognized that heterogeneity in the selective effect of TE insertional mutations may be necessary to maintain TEs in host populations (Charlesworth 1991). Previous models with fixed selective effects could not explain TE maintenance given reasonable parameter combinations calculated from observed data in *Drosophila* (Charlesworth and Langley 1989). Charlesworth’s deterministic model of both selected sites and neutral sites allows for the invasion of the population by TEs, but this results in cases with very high neutral copy number equilibriums, which, once again, conflicts with what is observed in *Drosophila*. Our simulations align with these results. We observe an accumulation of neutral TEs for most parameter combinations of transposition and insertion preference that is often larger ( > 100) than observed copy numbers in natural populations; however, as population size can fluctuate in our simulations, we find that these cases almost always result in host population extinction.

Our results suggest that aspects of TE biology unexplored in our models may be necessary for reaching high TE copy numbers. Charlesworth (1991) additionally considers a model where no transposition occurs from elements at neutral sites (modeling an insertion into centric heterochromatin which would be selectively neutral but silenced as a result) and found it relatively easy to find parameter combinations that result in much more realistic copy numbers for neutral and selected sites. Managed bursts of transposition may allow the copy number to grow and then allow selection purge deleterious insertions. Mechanisms that modulate transposition are varied and well-documented across a wide variety of TE families and among different species (Slotkin and Martienssen 2007; Fultz et al. 2015; Choi and Lee 2020). They can, however, these mechanisms can broadly be characterized as either auto-regulation or host repression. Auto-regulation occurs when TEs evolve self-regulatory mechanisms to modulate transposition depending on their own copy number. Studies in *Drosophila* found two mechanisms by which the mariner TE Mos1 regulates its own transposition, one of which reduces overall transpose activity as the number of active TE copies grows above some genomic optimal value (Lohe and Hartl 1996). Similarly, in *Saccharomyces* a retrotransposon-encoded restriction protein derived from the Ty1 element induces copy number control such that transposition is inhibited as the copy number increases (Saha et al. 2015). Host repression occurs when the host genome evolves to identify TEs and keep them from transposing. Chromatin modifications, germline silencing of TEs, and post-transcriptional silencing of TEs by RNA are all examples of ways with which host populations evolve to silence TE proliferation (see Slotkin and Martienssen 2007 for a review), and many of the genes involved in the latter are conserved across eukaryotes (Cerutti and Casas-Mollano 2006).

Transposition regulation occurs in varied ways, with both the TE family and the host genome potentially evolving to do so. It is unclear whether all TE families are capable of monitoring their genome-wide copy number and subsequently reducing transposition or whether it is an evolved trait given the right blend of biology and selective pressure. Similarly, while many species may already have evolved mechanisms for repressing TE proliferation, they may not have evolved the ability to recognize novel TEs. The piRNA pathway, which is a specific RNA silencing mechanism, leverages piRNAs to recognize targets and silence TEs (Brennecke et al. 2007; Tóth et al. 2016). piRNAs often originate from distinct piRNA clusters, which represent a few genomic loci that are highly enriched in transposable element sequences. It is hypothesized that once a novel TE inserts into this sequence it can then be targeted by the piRNA machinery to limit further transposition of the TE. As the copy number of the TE family increases, the chance of landing in a piRNA cluster increases as well. This suggests that host repression may also evolve in a manner that is dependent on the copy number of the TE family. Exploring the relationship between TE copy number and the evolution of TE regulation may be key to understanding how high TE copy numbers are achieved.

Our models assume the introduction of a single TE into a naïve population in a mechanism similar to horizontal transfer. However, TEs could be introduced at higher copy numbers into populations via hybridization and subsequent introgression. Barbara McClintock hypothesized that hybridization between different populations or species could act as a “genomic shock”, as TEs began to proliferate again (McClintock 1984). This shock could be detrimental as hybrids could exhibit reduced fitness as a consequence of TE proliferation. One of the most well characterized examples of this is the hybrid dysgenesis system in *Drosophila melanogaster*, where intraspecific crosses between strains carrying the P-element transposon to those without it result in sterile offspring (Kidwell et al. 1977; Bingham et al. 1982; Kidwell 1983; Labrador et al. 1999; Kelleher et al. 2012).

An important feature of TE biology that is absent in our model is the adaptive potential of TEs (Li et al. 2018). Inserted TEs can alter or act as novel coding sequences in the genome altering the expression levels of genes (Hoen and Bureau 2015; Joly-Lopez et al. 2016). Similarly, the epigenetic modifications that result from TE insertions may also change gene expression patterns (Lisch 2013; Stuart et al. 2016). In maize, the increased apical dominance compared to its progenitor, teosinte, is partially explained by a TE insertion upstream of the domestication gene *tb1* that now acts as an enhancer of gene expression (Studer et al. 2011). TEs may consequently contribute in important ways to rapid adaptation by inducing gene modifications, altering gene expression, or exaptation to create novel genes (Oliver et al. 2013; Schrader et al. 2014; Stapley et al. 2015). We hypothesize that the addition of adaptive TE insertions may result in cyclic fitness trajectories. An adaptive TE will sweep through the population boosting mean population fitness, temporarily overcoming the growing load of deleterious TEs. Once the adaptive TE fixes, the accumulation of deleterious TEs will continue increasing and reducing population fitness again. How readily adaptive TEs appear will consequently determine whether populations can avoid entering into an extinction vortex.

In total, our results question the assumption that evolving a preference for neutral TE insertions is protective. We find that this preference, while useful in terms of increasing copy number, ultimately leads to population extinction and the elimination of the TE family. It also raises several questions about how high TE copy number can be achieved without resulting in host population extinction. Dual survival, in our model, was predominantly characterized by low copy number, and, in natural populations, many TE families exhibit single or low copy numbers as well (Stitzer et al. 2021; Stritt et al. 2021). Extinction eliminates both the host population and the TE family. If TEs drive host lineages extinct before becoming prevalent in other genetic backgrounds, the TE can fail to invade (we see a failure to invade with modest selection coefficients, low neutral insertion preferences, and low recombination rates (Fig 2A, 2C, and 2D). More often, highly proliferative TEs can drive entire host populations extinct well after the TE has established. This may shape the evolution of both the host and the TE in unexplored ways. The interplay between extinction and the pressure to replicate or reproduce will ultimately determine not only how TE copy number changes over time but also how hosts respond to a TE invasion.

