## Supplemental Data for "Critical Role of Insertion Preference for Invasion Trajectory of Transposons"

**Life Cycle, Population Growth, and Selection** – The generational life cycle of the non-Wright-Fisher framework in SLiM is detailed in the manual (Haller and Messer 2016) and can be summarized as follows: generation of offspring, execution of early() events to adjust either individual genomes or the population, calculation of individual fitness, viability selection on present individuals, and then execution of late() events which provide another opportunity to adjust genomes or the population before the cycle loops.

We initialize the population with $K$ = 1000 individuals, with $K$ acting as the hard carrying capacity for the population. The generational life cycle in non-Wright-Fisher models in SLiM starts with the creation of offspring. We generate twice the current population size (2$N$) offspring each generation with the two parents for each individual chosen at random. Without this adjustment, the population rapidly experiences extinction as even the loss of a single individual can never be recovered (i.e., the loss of a single individual per generation due to selection would drive the population extinct in 1000 generations). Consequently, we generate twice as many offspring and then cull to the carrying capacity.

Non-Wright-Fisher models employ viability selection, which is based on a probabilistic calculation of fitness where values less than or equal to 0 ensure death and values equal to 1 ensure survival. We enforce discrete generations by setting the fitness of all parents to 0 before viability selection occurs. Similarly, we cull excess offspring if $2N > K$ by randomly selecting individuals (agnostic to their final fitness) and setting their fitness to 0 prior to selection. Due to the generational life cycle of SLiM, culling requires us to set the selected individuals fitness to 0 prior to viability selection. Viability selection will then automatically remove parents and culled individuals, while the remaining individuals probabilistically survive with higher fitness individuals having a greater chance of survival. Transposition then occurs in the remaining individuals who survived viability selection. Since fitness is evaluated only during viability selection, new or lost elements do not influence the probability of an individual being chosen as a parent in the next generation. They will, however, be inherited by any potential offspring, and, consequently, influence their offspring's ability to survive. We then loop back to the start of the generational cycle with the surviving offspring forming the new parental pool.

**Model Outcomes** – For each distinct parameter combination, we track the mean fitness and size of the population as well as the mean copy number of the TE family and mean frequency of a TE in the family stratified by their fitness effects over time. We consider 3 endpoints for a replicate simulation run of a given parameter set.

*Outcome 1, TE Loss* – Loss of the TE family from the host population occurs if the mean copy number of autonomous TEs is zero. No further transposition can occur, and we simply track which generation the TE family was lost in.

*Outcome 2, Population Extinction* – If the population size hits zero, we record that replicate as resulting in a population extinction event and output the relevant trajectories (population fitness, size, mean TE copy number, and mean TE frequency over time). population extinction occurs when no individuals in the population survive after viability selection.

*Outcome 3*, Dual Survival – If neither of these options occur, then both the TE family and the host population have survived to the final generation (capped at 50,000). We output the relevant trajectories (same as though outputted in population extinction) for that simulation run.

**Models, Parameters, and Replicates** – We consider 5 distinct models that varied either genome architecture or specific aspects of TE biology detailed above (Table 1). When describing our results, we use the term TE family to refer to a specific parameter combination of transposition probability (*teJumpP*) and insertion preference (*neutP*). We explore the following sets of values for our parameters: *teJumpP* = [$1 \times 10^{-4}$, $2.5 \times 10^{-4}$, $5 \times 10^{-4}$, $7.5 \times 10^{-4}$, $1 \times 10^{-3}$, $2.5 \times 10^{-3}$, $5 \times 10^{-3}$, $7.5 \times 10^{-3}$, $1 \times 10^{-2}$, $2.5 \times 10^{-2}$, $5 \times 10^{-2}$, $7.5 \times 10^{-2}$, $1 \times 10^{-1}$] (13 values) and *neutP* = [0.010, 0.025, 0.050, 0.075, 0.10, 0.25, 0.50, 0.75, 0.90, 0.925, 0.950, 0.975, 0.99] (13 values) for each model, resulting in a total of 169 distinct parameter combinations per model type. Each distinct parameter combination is used for a specific simulation run of a model. We initialize the host genome, TE family, and population and consider 3 endpoints (TE loss, population extinction, or dual survival) for a replicate of a given parameter set. When one of these outcomes occur, we record the relevant trajectories and end state before looping back to the initialized state. The simulation run is finally complete (i.e., we no longer loop back to the beginning point) if we record either a combined total of 100 population extinction and dual survival events or $1 \times 10^{6}$ TE loss events. Certain parameter combinations resulted in dramatically longer run times, due to SLiM executing operations over a large number of TEs for many generations. For these combinations, we scaled down the outcome by an order of magnitude (i.e., 10 population extinction and dual survival outcomes to 1 x $10^5$ TE Loss outcomes) and parallelized runs if necessary (see Supplementary Table 1 for parameter combinations that were scaled). If runs needed to be parallelized, we simply scaled down by another order of magnitude (i.e., 1 population extinction and dual survival outcomes to 1 x $10^4$ TE Loss outcomes) and aggregated 10 runs into a single replicate. In all figures presented, we mark these combinations to highlight this change. Scripts for all models can be found on GitHub – see Data accessibility statement

**Table S1. Scaled Down Parameter Combinations**

See Attached CSV

**Table S2. TE Loss Outcomes Across Models**

| Model | Minimum TE Loss Proportion | Percent of TE Loss Outcomes Within 10 Generations | Percent of TE Loss Outcomes Within 50 Generations | Percent of TE Loss Outcomes Within 500 Generations |
|---|---|---|---|---|
| Model 1 – Single Chromosome | 87.67% | 68.64% | 95.68 | 99.67% |
| Model 2 – Single Chromosome With Excision | 90.19% | 68.69% | 95.72% | 99.67% |
| Model 3 – Single Chromosome With Non-Autonomous Elements | 93.71% | 68.64% | 95.85% | 99.78% |
| Model 4 – Single Chromosome With Excision & Non-Autonomous Elements | 94.76% | 68.85% | 95.97% | 99.75% |
| Model 5 – Five Chromosome | 84.66% | 68.83% | 95.78% | 99.62% |

Column 1 - Model Type
Column 2 - The minimum TE Loss proportion across the entirety of the parameter space, the specific parameter combination is listed below in parentheses
Column 3 - Across the entire parameter space, the percentage of TE loss outcomes that occurred in less than 10 Generations
Column 4 - Across the entire parameter space, the percentage of TE loss outcomes that occurred in less than 50 Generations
Column 5 - Across the entire parameter space, the percentage of TE loss outcomes that occurred in less than 500 Generations

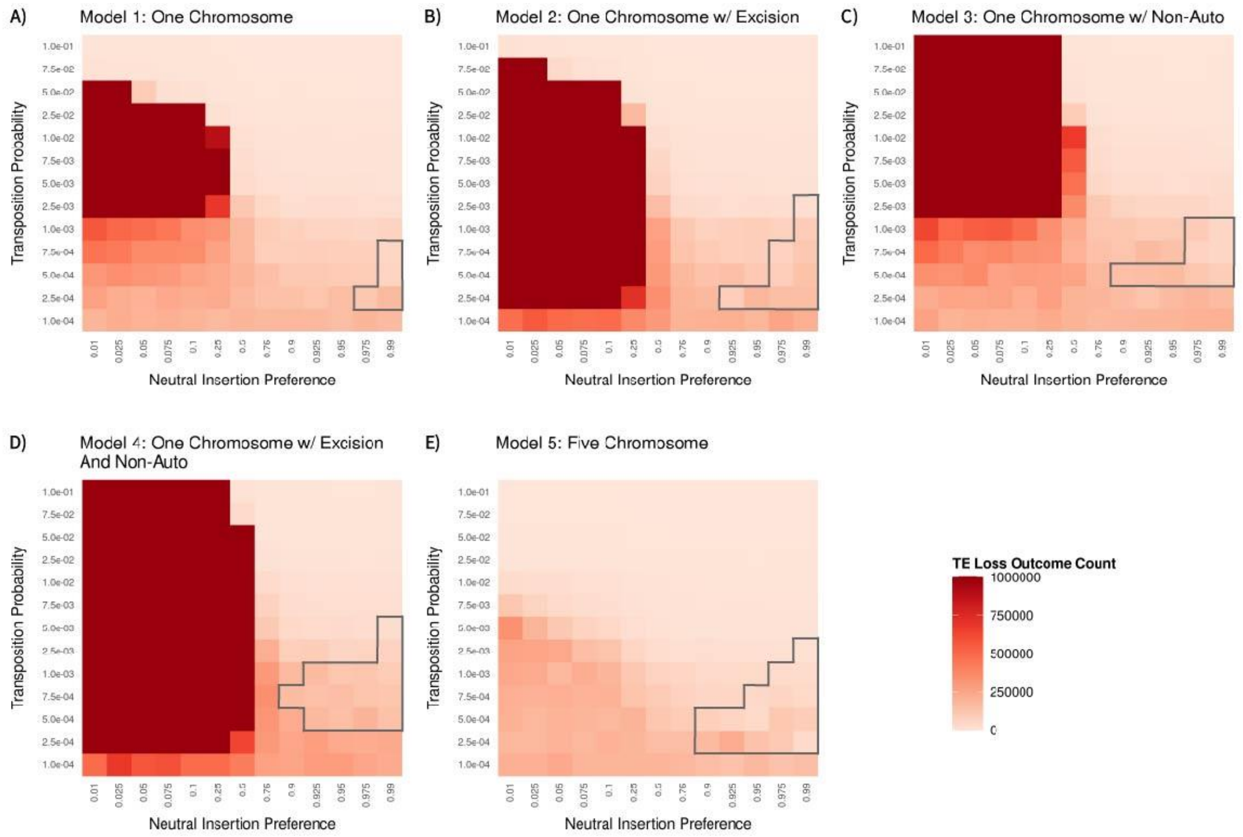


**Figure S1 TE Loss Outcomes Across Our Parameter Space by Model Type.** For each heatmap, rows represent the transposition probability with increasing probabilities as you move upwards, and columns represent the neutral insertion preference with increasing preferences as you move to the right. Darker colors indicate more TE Loss outcomes. Cells indicated with grey markers represent parameter combinations that, due to computational constraints, had reduced limits detailed in the methods. The heatmaps show the number of TE Loss Outcomes for our (A) One Chromosome Model, (B) One Chromosome Model with Excision, (C) One Chromosome Model With Non-Autonomous Elements, (D) One Chromosome Model with Excision and Non-Autonomous, and (E) Five Chromosome Model.

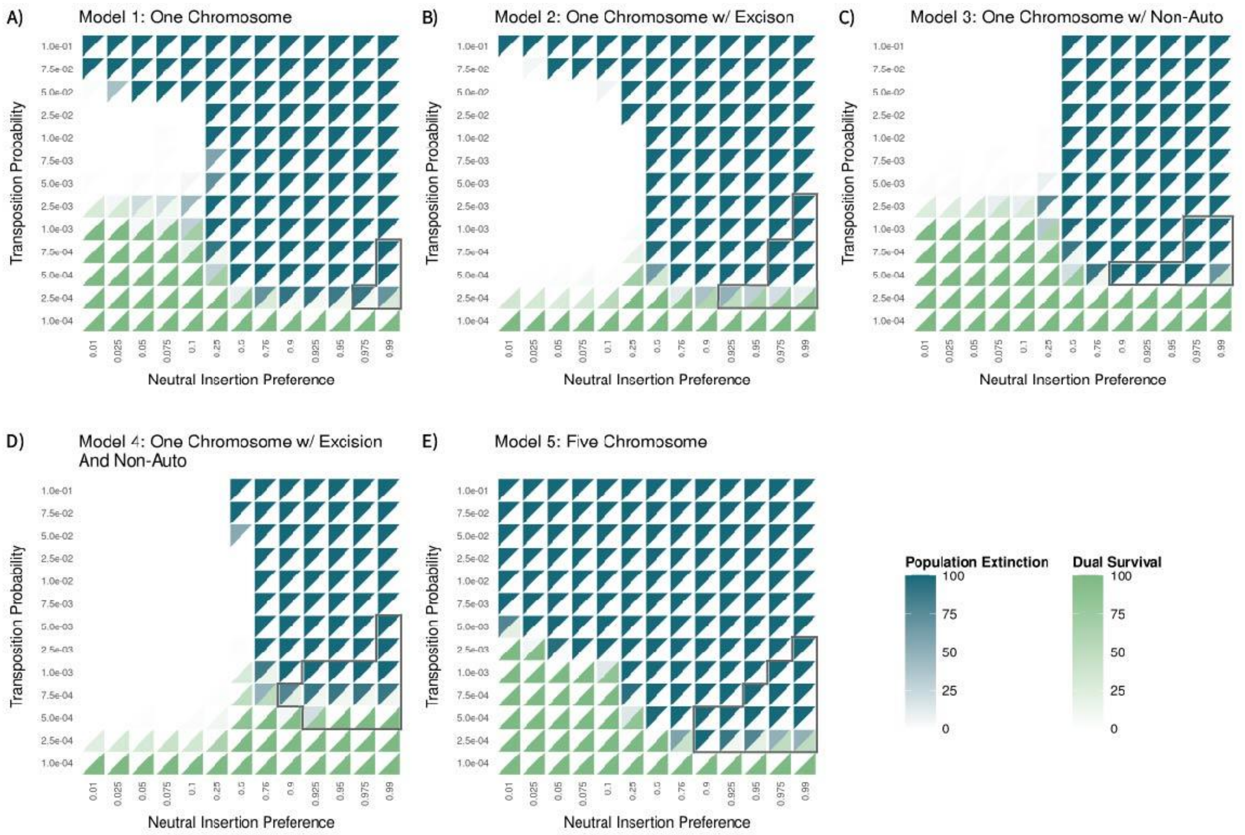


**Figure S2 Population Extinction and Dual Survival Outcomes Across Our Parameter Space by Model Type.** For each heatmap, rows represent the transposition probability with increasing probabilities as you move upwards, and columns represent the neutral insertion preference with increasing preferences as you move to the right. Heatmaps are colored with both the number of TE introductions that resulted in population extinction (upper triangle in blue) or dual survival (lower triangle in green) with darker colors representing higher numbers. Cells indicated with grey markers represent parameter combinations that, due to computational constraints, had reduced limits detailed in the methods. The heatmaps show the number of outcomes for our (A) One Chromosome Model, (B) One Chromosome Model with Excision, (C) One Chromosome Model With Non-Autonomous Elements, (D) One Chromosome Model with Excision and Non-Autonomous, and (E) Five Chromosome Model.

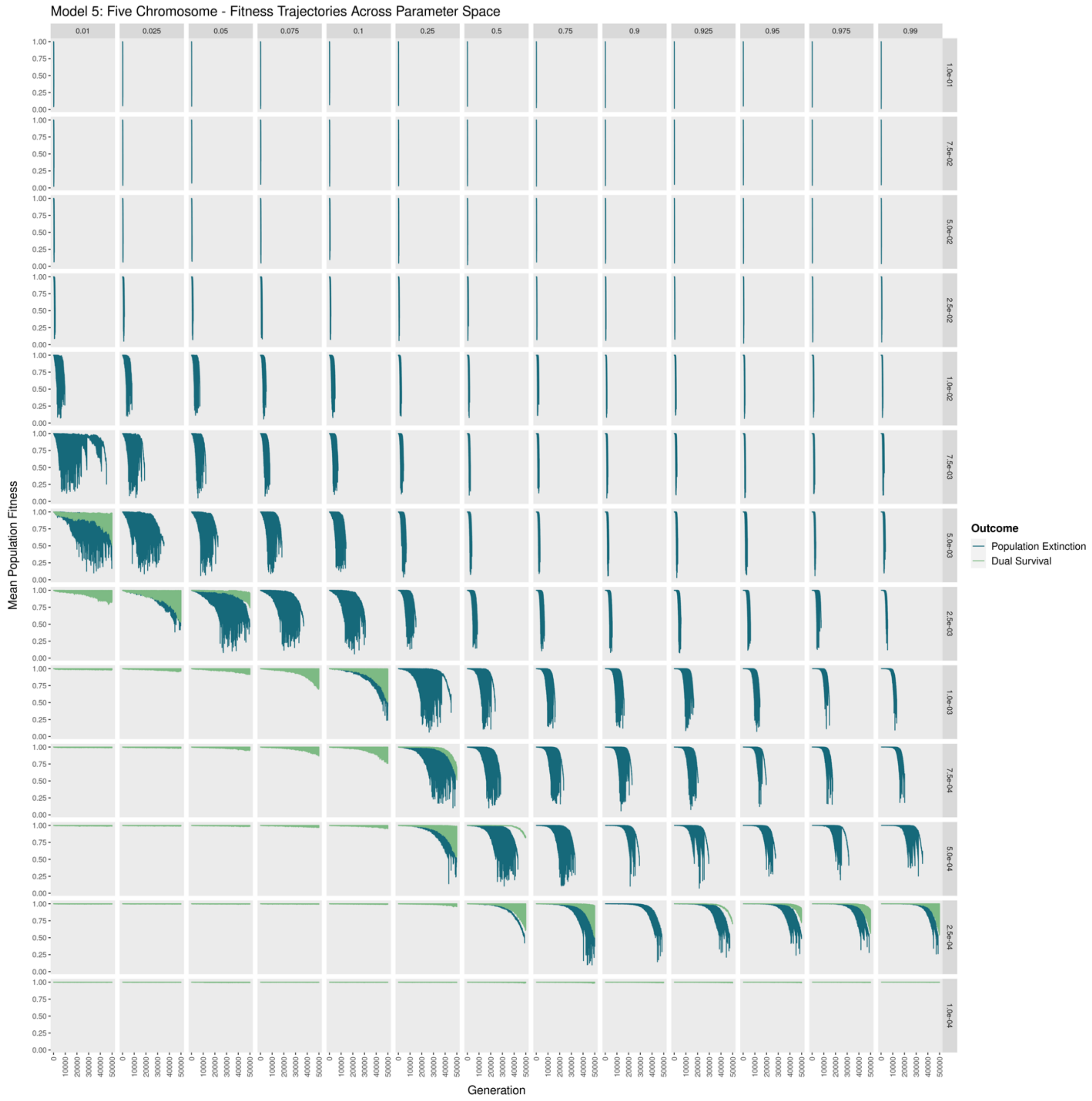


**Figure S3 Population Fitness Trajectories Over Time Across Our Parameter Space For Our Five Chromosome Model**. We compare mean population fitness trajectories across our parameter space, where rows represent the transposition probability, with greater probabilities on the top, and columns represent the insertion preference, with stronger preferences for neutral insertion sites on the right. Each line represents the mean population fitness trajectory for a given replicate that resulted in either population extinction or dual survival. Population extinction trajectories are colored in dark blue, while dual survival outcomes are colored in green.

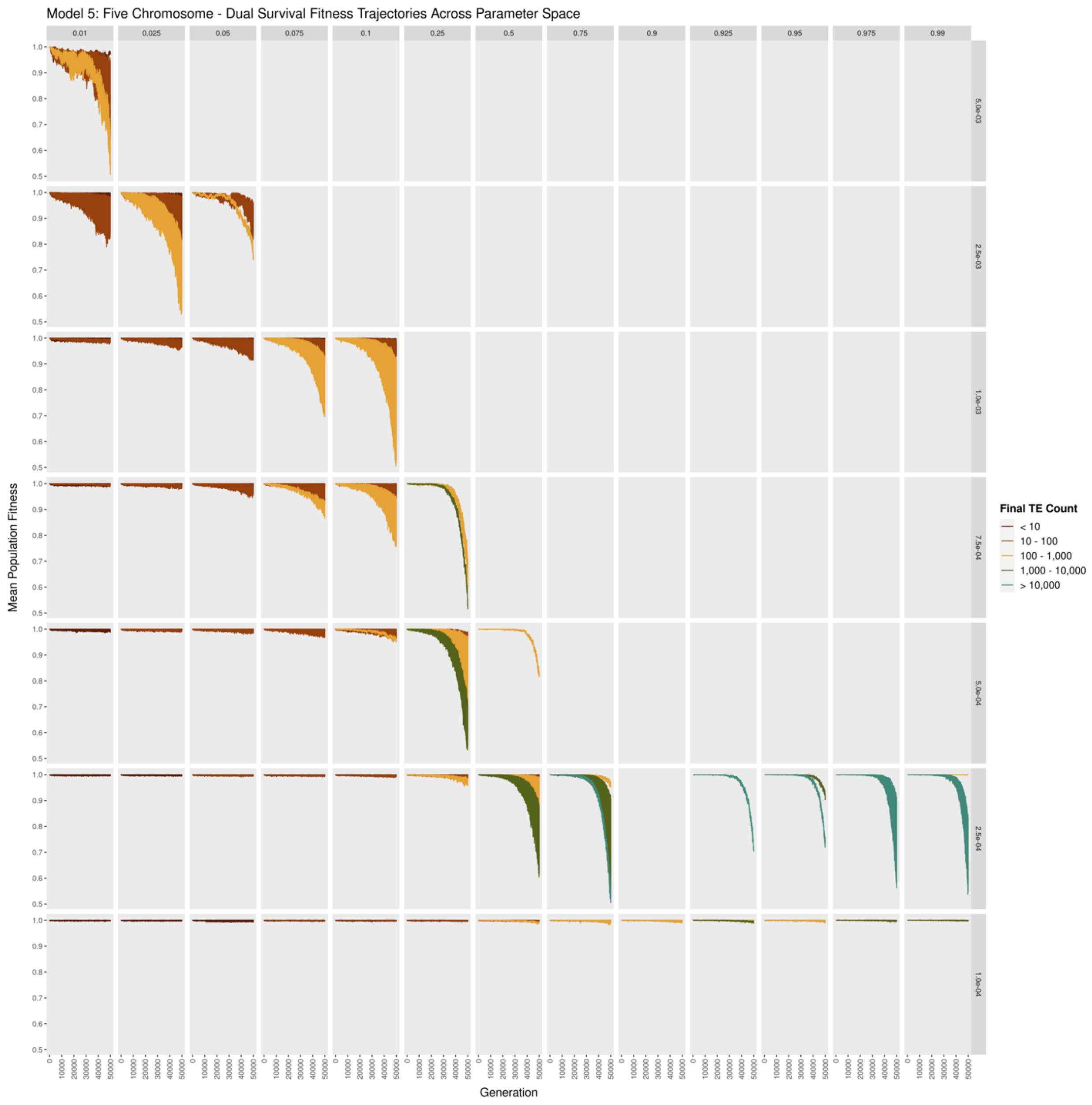


**Figure S4 Population Fitness Trajectories Over Time Across Our Parameter Space For Our Five Chromosome Model for Dual Survival Outcomes**. We compare mean population fitness trajectories across our parameter space, where rows represent the transposition probability, with greater probabilities on the top, and columns represent the insertion preference, with stronger preferences for neutral insertion sites on the right. Each line represents the mean population fitness trajectory for a given replicate that resulted in either population extinction or dual survival. We color trajectories by the final mean TE copy number for that replicate to show how many of our high TE copy number dual survival outcomes exhibit fitness trajectories similar to population extinction.
